## Supplemental Figures for "Deep mutational scanning reveals a tight correlation between protein degradation and toxicity of thousands of non-native aspartoacylase protein variants"

*Supplemental material*


**Figure S1**, *No detectable endogenous ASPA in HEK293T cells.* p.2
**Figure S2**, *Abundance score correlations between independent repeats.* p.3
**Figure S3**, *Zoom in on the $Zn^{2+}$ coordinating residues and β-strand at 150-160.* p.4
**Figure S4**, *Many low abundance variants are buried.* p.5
**Figure S5**, *Comparisons of the abundance map with the Rosetta stability predictions.* p.6
**Figure S6**, *Comparisons of the abundance map with the in silico predictions.* p.7
**Figure S7**, *Inherent degrons are buried in the ASPA structure.* p.8
**Figure S8**, *The screen for toxic ASPA variants.* p.9
**Figure S9**, *Toxicity score correlations between independent repeats.* p.10
**Figure S10**, *Toxicity score distribution.* p.11
**Figure S11**, *ASPA toxicity scores and abundance determined in low throughput.* p.12
**Figure S12**, *Comparisons of the mutational maps.* p.13
**Figure S13**, *Toxic variants are typically buried.* p.14
**Figure S14**, *Benign variants are not toxic.* p.15
**Figure S15**, *Toxic variants activate a stress response leading to HSP70 induction.* p.16

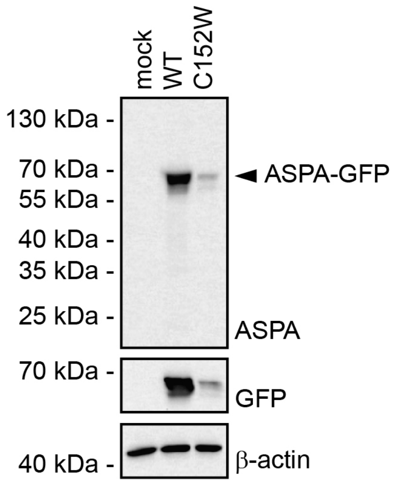


**Figure S1** – *No detectable endogenous ASPA in HEK293T cells.*
The level of endogenous and recombinant ASPA in the HEK293T landing pad cell line was analyzed was by SDS-PAGE and western blotting using antibodies to ASPA and GFP. β-actin served as a loading control.

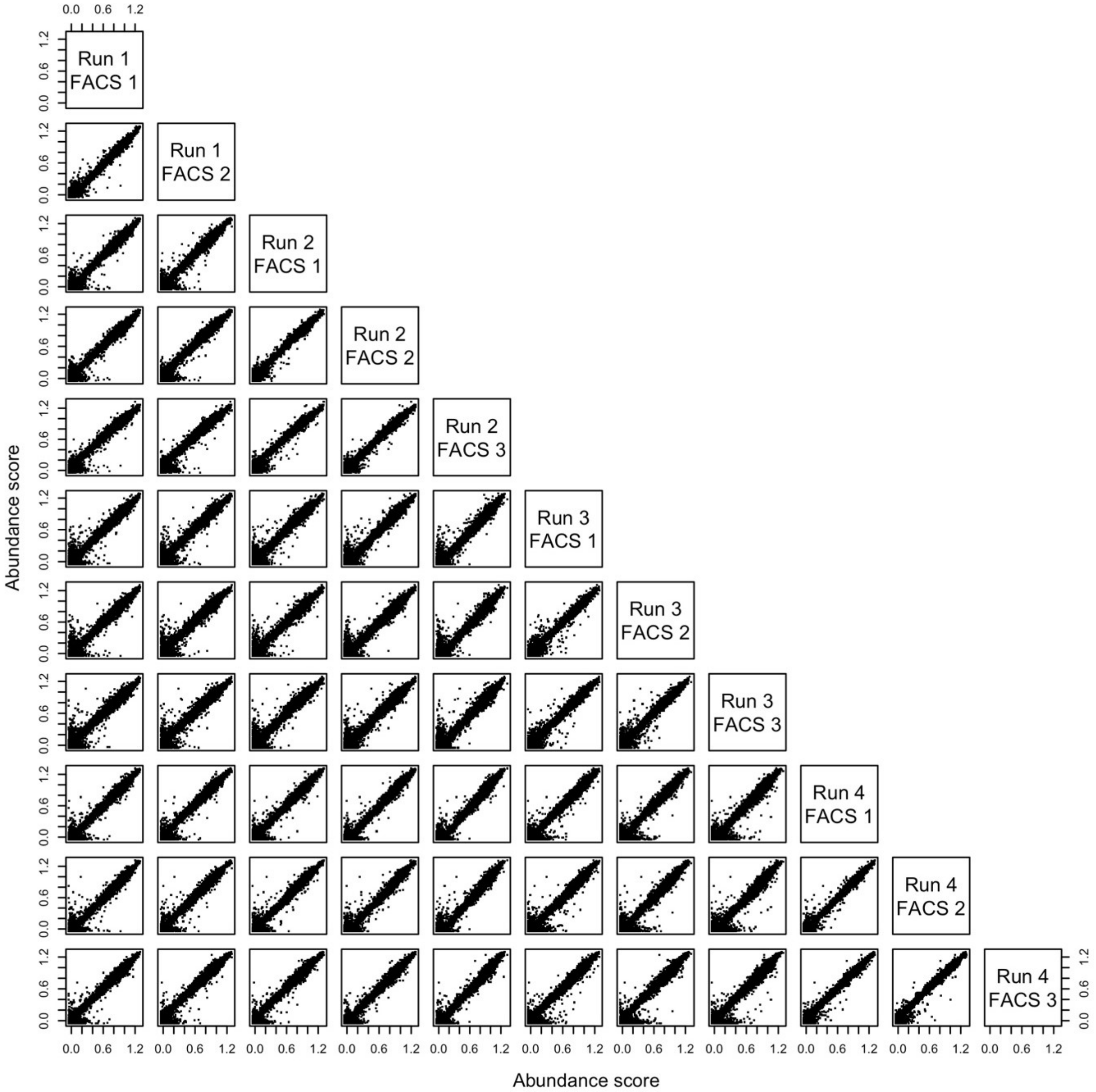


**Figure S2** – *Abundance score correlations between independent repeats.*
Correlation of single amino acid variant scores between all four biological replicates (Run 1-4) each with three FACS replicates except for B1 which only have two FACS replica. All Pearson correlations are in the range 0.98 to 0.99.

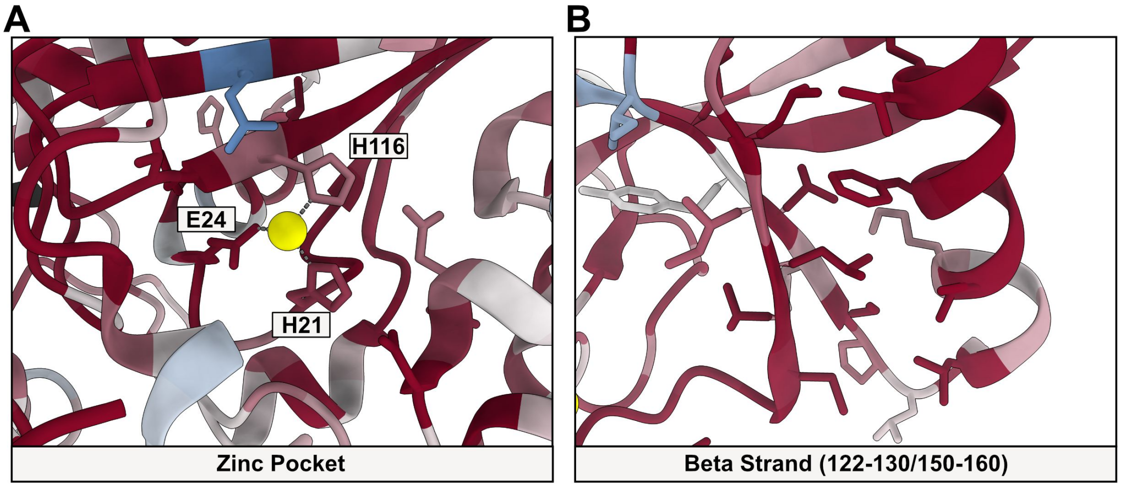


**Figure S3** – *The $Zn^{2+}$ coordinating residues and β-strand at 150-160.*
(A) The $Zn^{2+}$ coordinating residues in ASPA. The structure is colored based on the median abundance score per residue. (B) The alternating pattern of low and high abundance variants in the β-strand at position 150-160 is shown. The structure is colored based on the median abundance score per residue.

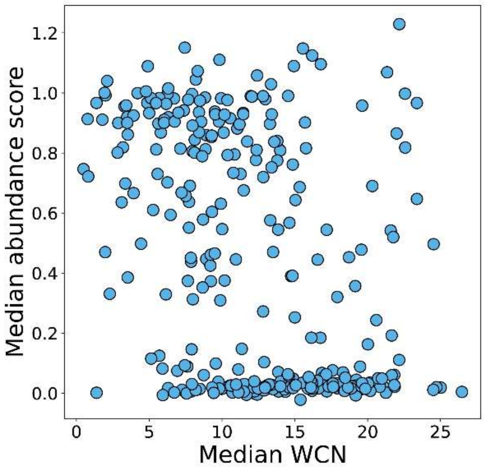


**Figure S4** – *Many low abundance variants are buried.*
Plot of the median abundance score vs. the weighted contact number (WCN).

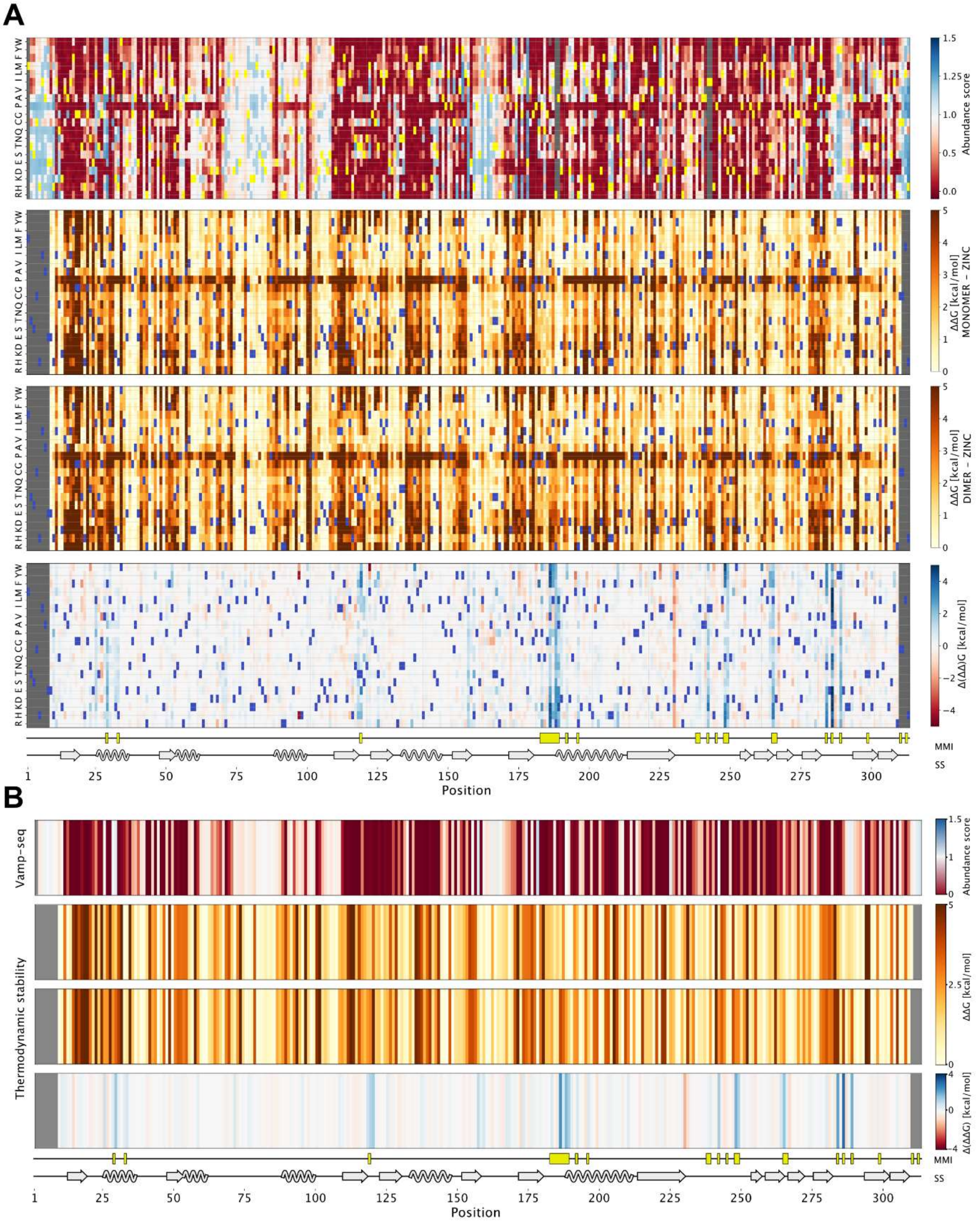


**Figure S5** – *Comparisons of the abundance map with the Rosetta stability predictions.*
(A) Side-by-side comparisons of the ASPA abundance map with the Rosetta stability maps for the ASPA dimer (with zinc) and ASPA monomer (without zinc). The difference in predicted stability (ΔΔΔG) between the ASPA monomer and dimer is also included (lower panel). The secondary structure (SS) elements and monomer-monomer interface (MMI) positions are included for comparison. (B) Comparisons of the ASPA median abundances per residue with the corresponding Rosetta stability predictions for the ASPA dimer and monomer. The difference in predicted stability (ΔΔΔG) between the ASPA monomer and dimer is also included (lower panel). The secondary structure (SS) elements and monomer-monomer interface (MMI) positions are included for comparison.

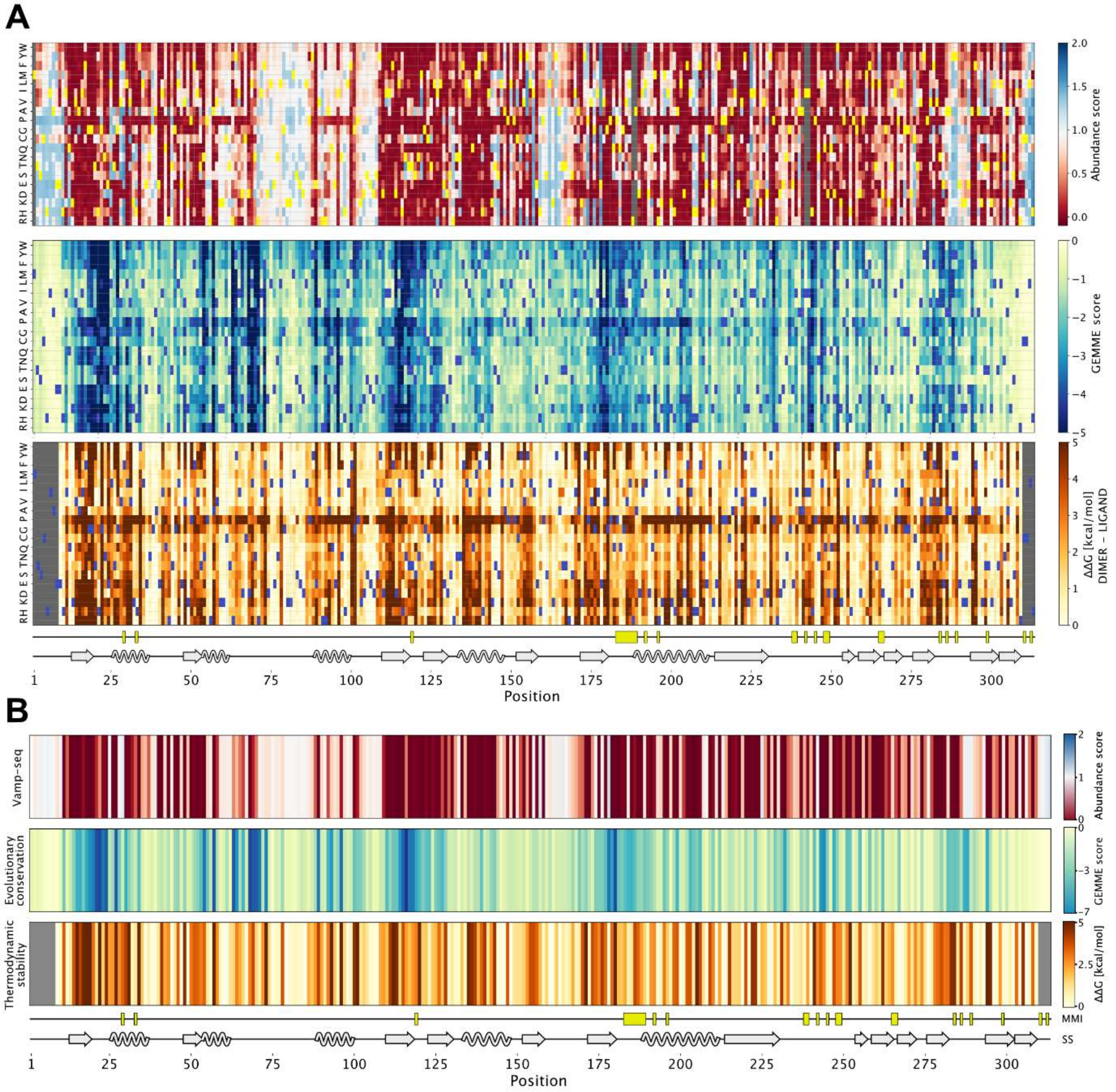


**Figure S6** – *Comparisons of the abundance map with the in silico predictions.*
(A) Side-by-side comparisons of the ASPA abundance map with the GEMME (ΔΔE) conservation and Rosetta stability maps for the ASPA dimer. The secondary structure (SS) elements and monomer-monomer interface (MMI) positions are included for comparison. (B) Comparisons of the ASPA median abundances per residue with the corresponding residue median GEMME and median Rosetta stability predictions for the ASPA dimer. The secondary structure (SS) elements and monomer-monomer interface (MMI) positions are included for comparison.

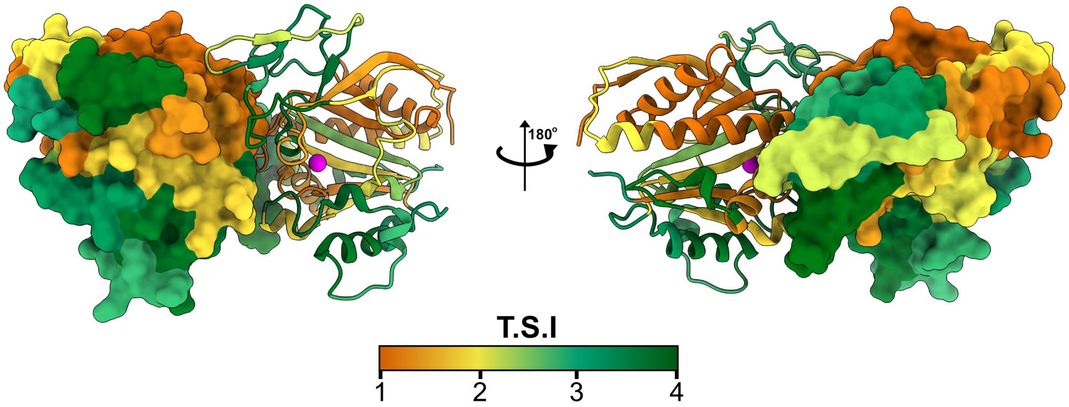


**Figure S7** – *Inherent degrons are buried in the ASPA structure.*
The ASPA structure colored by TSI. Note that regions with low stability (degrons) are mostly buried in the structure.

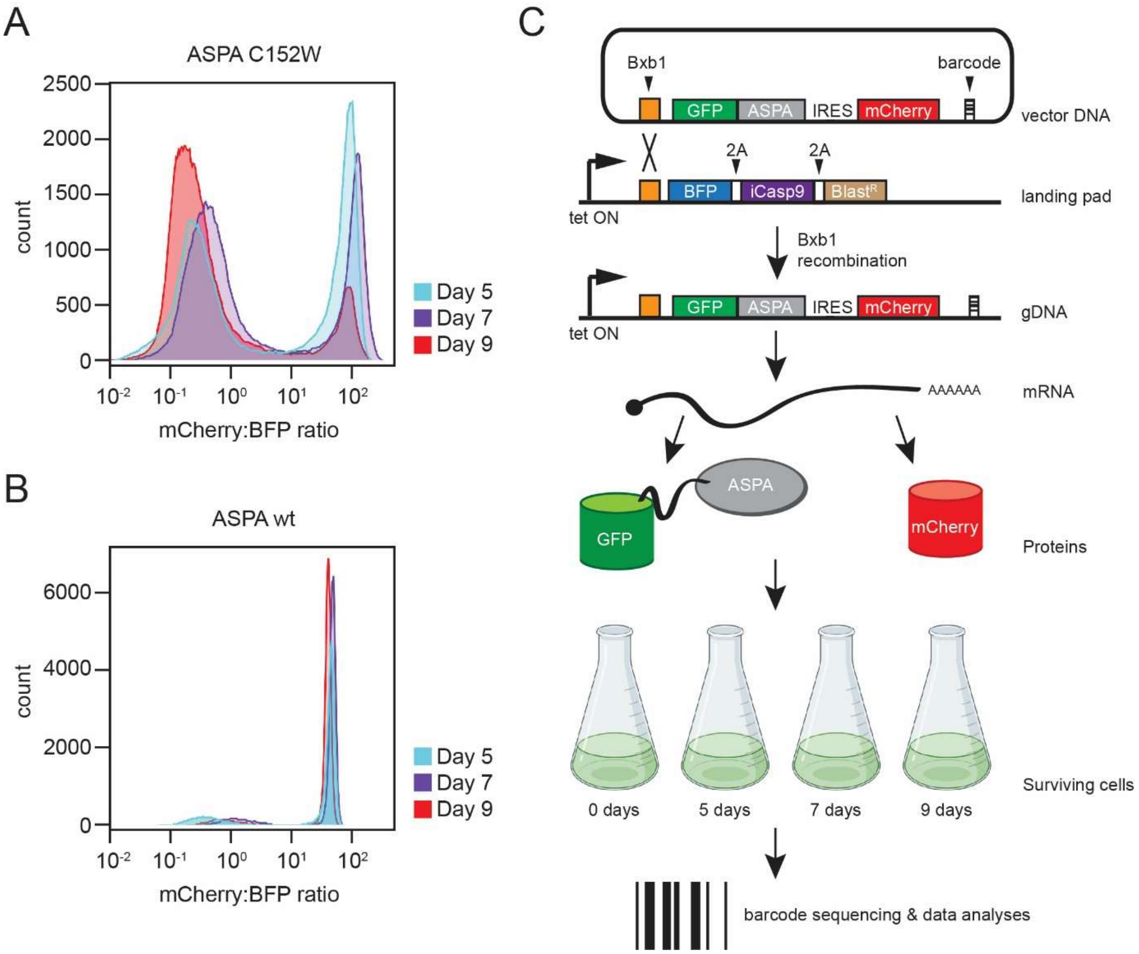


**Figure S8** – *The screen for toxic ASPA variants.* (A) Histogram of the mCherry:BFP ratio based on flow cytometry ASPA C152W after 5 (cyan), 7 (purple) and 9 (red) days of expression with doxycyclin. Note that with time the number of mCherry-expressing (ASPA C152W positive) cells declines, whereas the number of non-recombined (BFP positive) cells increase. (B) Histogram of the mCherry:BFP ratio based on flow cytometry wild-type ASPA after 5 (cyan), 7 (purple) and 9 (red) days of expression with doxycyclin. (C) Schematic representation of the expression and screening system. HEK293T cells carrying a landing pad for Bxb1-catalyzed site-specific integration are transfected with the expression vector and a Bxb1 expression plasmid (not shown). Upon integration at the landing pad locus, the BFP-iCasp9-Blast$^R$ gene is displaced downstream, and the cells therefore become resistant to AP1903, while GFP-ASPA and mCherry is expressed from the tetracyclin/doxycyclin regulated promoter. The same mRNA leads to both GFP-ASPA and mCherry protein production. Without flow sorting, cells were harvested after 0, 5, 7 and 9 days culturing with doxycyclin. Surviving variants were identified by sequencing the barcodes.

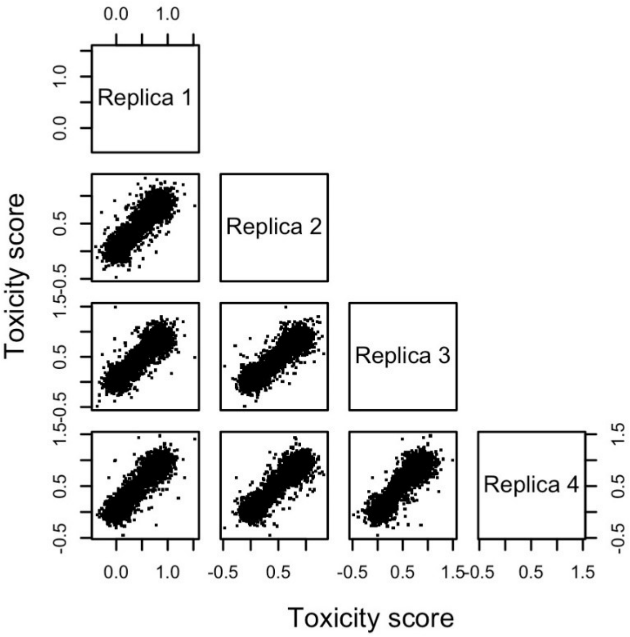


**Figure S9** – *Toxicity score correlations between independent repeats.*
Correlation of single amino acid variant scores between all four biological replicates. All Pearson correlations are in the range 0.93 to 0.94.

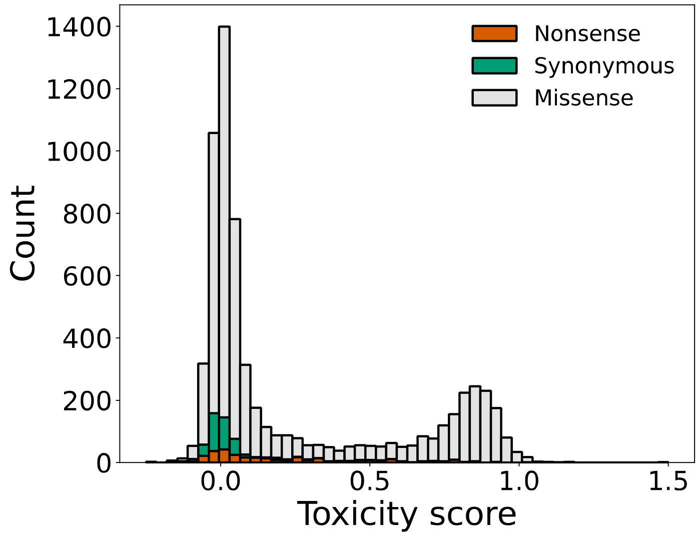


**Figure S10** – *Toxicity score distribution.*
Stacked histogram of toxicity scores for ASPA nonsense (red), synonymous (green) and missense (gray) variants. The missense variant toxicity score distribution is bimodal with a large peak around 0 originating from non-toxic variants and a smaller peak of toxic variants with toxicity scores close to 1.

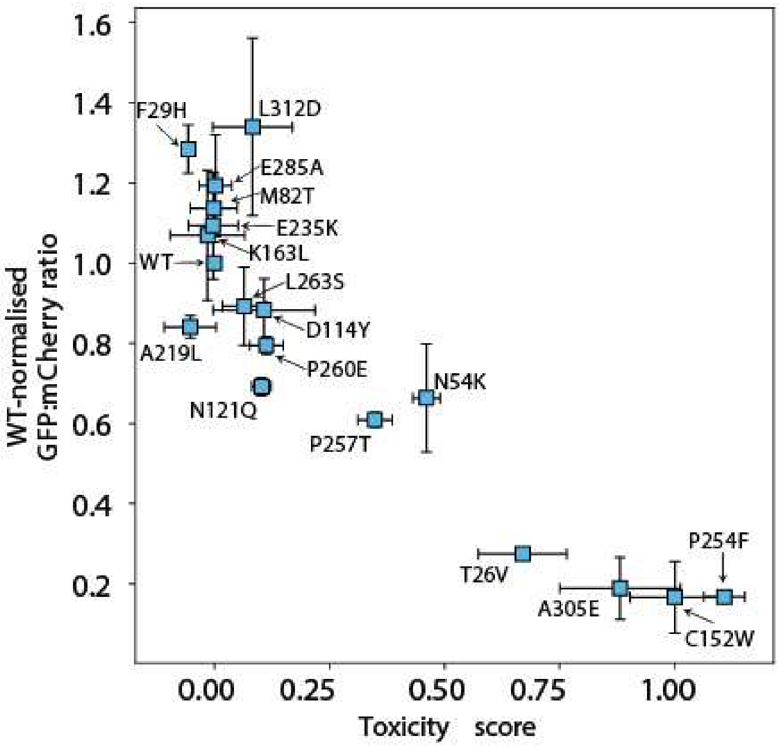


**Figure S11** –*ASPA toxicity scores and abundance determined in low throughput.*
To compare toxicity and abundance, 17 ASPA variants and wild-type ASPA were analyzed one-by-one by flow cytometry in low throughout. The abundance scores determined in low-throughput (y-axis) correlate with the toxicity scores determined from the screen (x-axis).

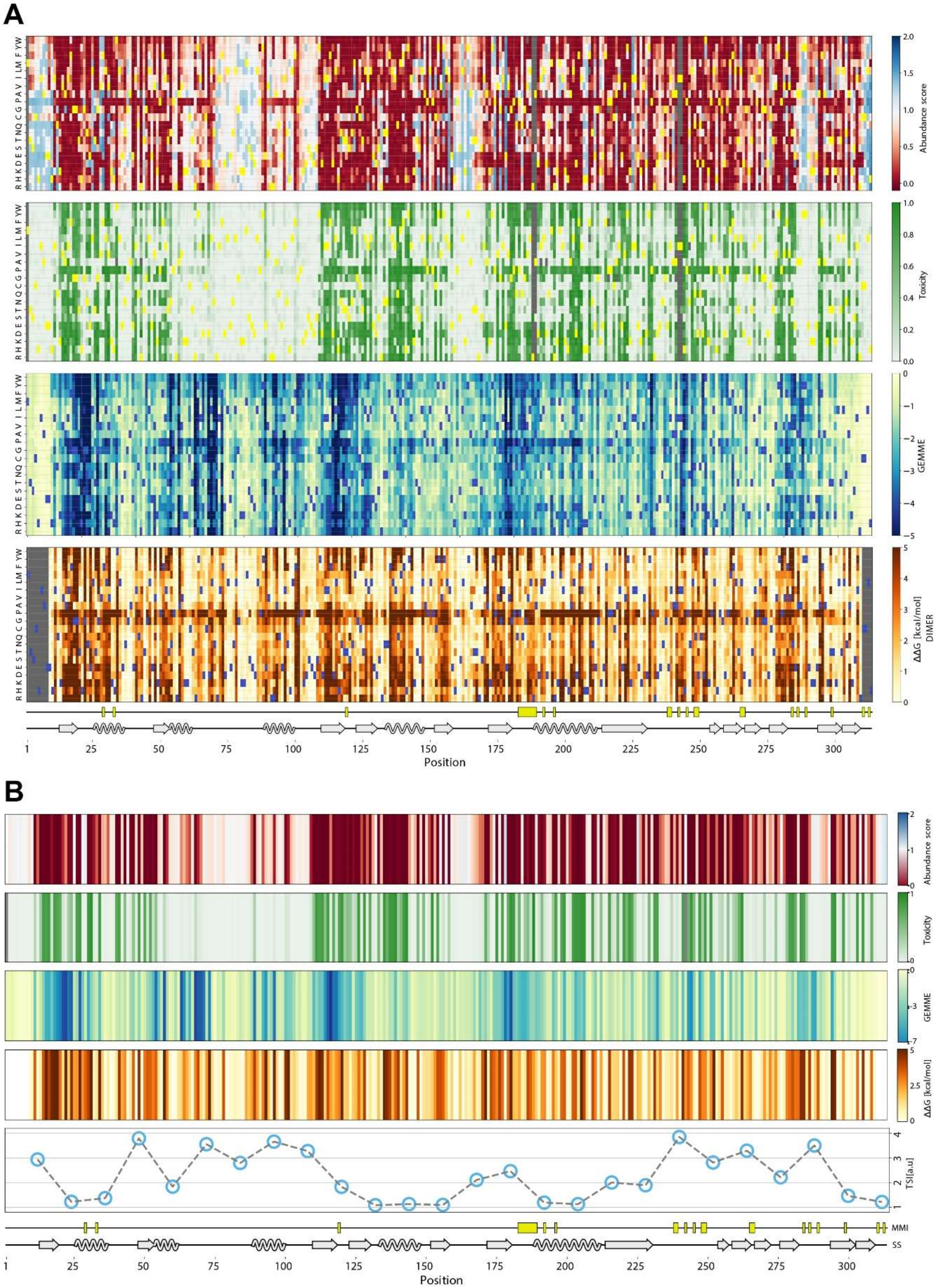


**Figure S12** – *Comparisons of the mutational maps.*

(A) Side-by-side comparisons of the ASPA abundance map, the toxicity map, the GEMME conservation scores and Rosetta predicted change in thermodynamic stability (ΔΔG) for the ASPA dimer. The secondary structure (SS) elements and monomer-monomer interface (MMI) positions are included for comparison. (B) Comparisons of the ASPA median abundances and toxicity per residue with the corresponding residue median GEMME and median Rosetta stability predictions for the ASPA dimer. The tile stability index (TSI) is plotted for the ASPA tiles for comparison.

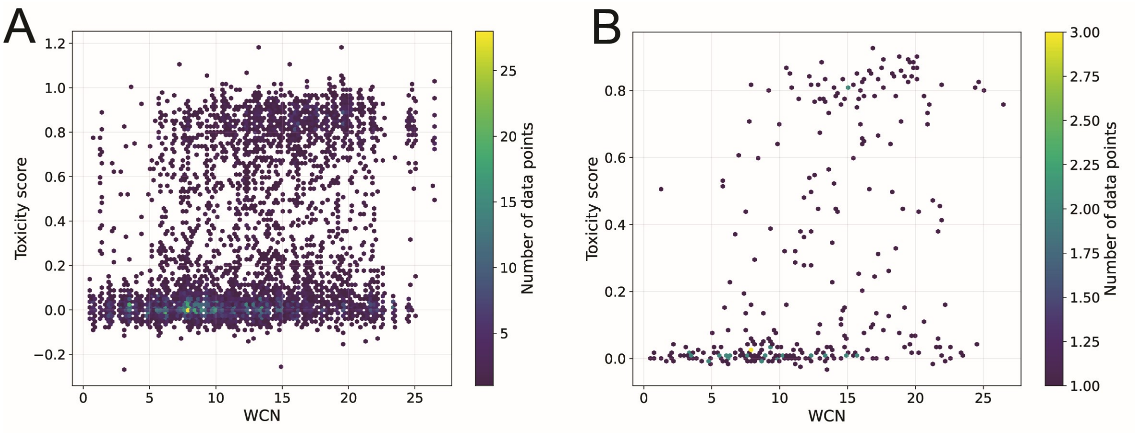


**Figure S13** – *Toxic variants are typically buried.*
Plots of the (A) toxicity scores and (B) median toxicity scores vs. the weighted contact number (WCN).

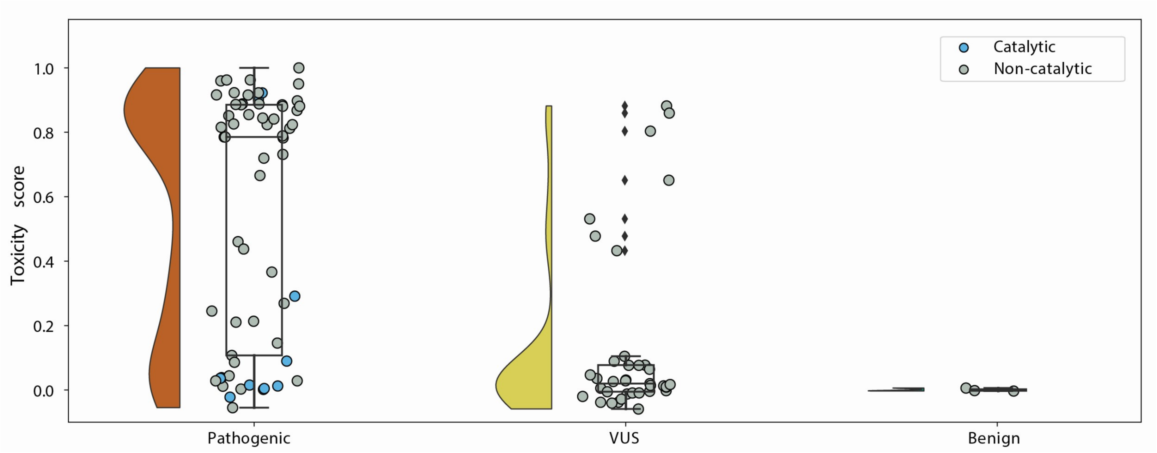


**Figure S14** – *Benign variants are not toxic.*

Comparisons of the toxicity scores for ASPA missense variants listed in Supplemental File 1 as pathogenic (red), variants of uncertain significance (VUS) (yellow) and benign (green) are shown as raincloud plots. Residues in or near the ASPA active site have been marked (blue). Note that many pathogenic and some VUS variants are toxic. Many of the non-toxic pathogenic variants are located near the active site (catalytic, blue).

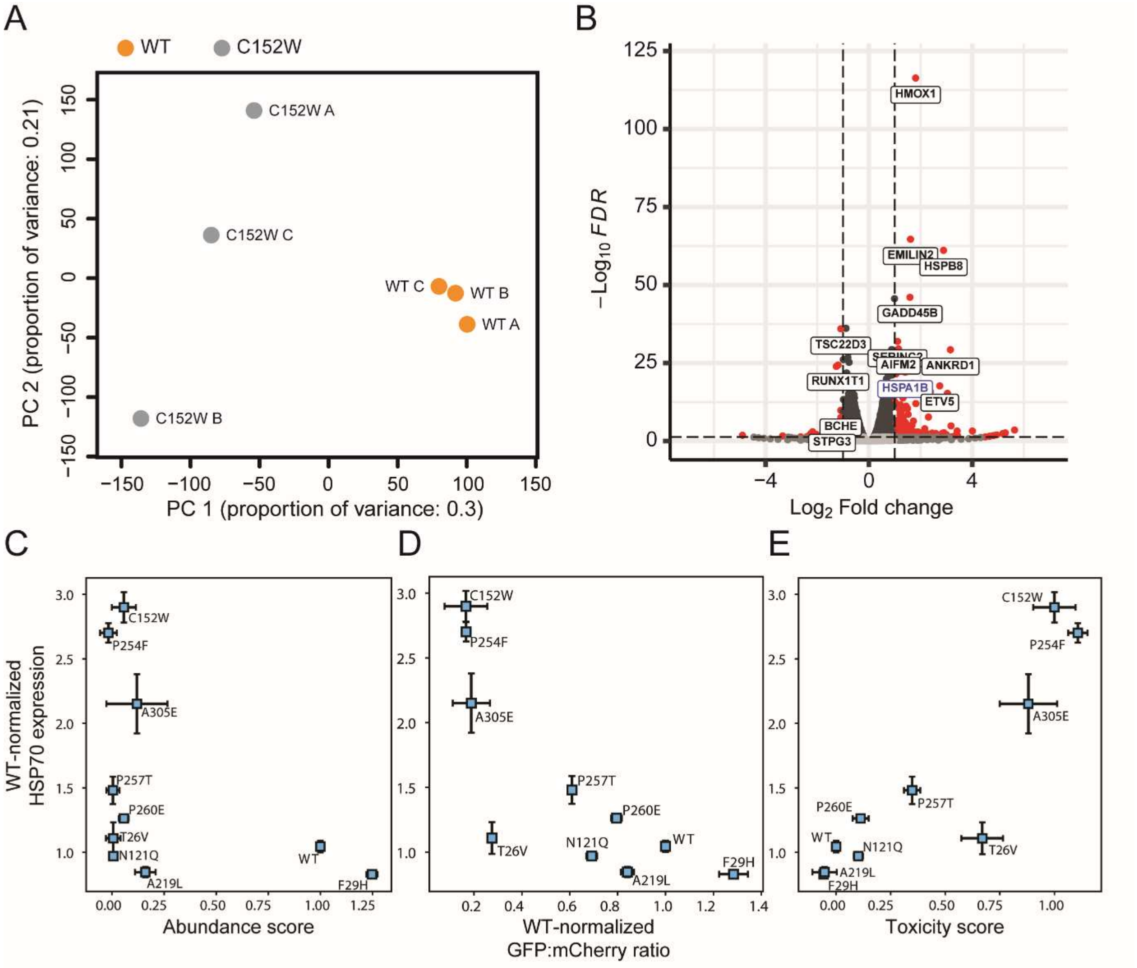


**Figure S15** – *Toxic variants activate a stress response leading to HSP70 induction.*
(A) Principal component (PC) analysis (PCA) of RNA sequencing on cells expressing wild-type (WT) ASPA (orange) or C152W (grey). The three independent repeats are marked A, B and C. (B) Differentially expressed genes between WT and C152W cells presented as a volcano plot. Genes where the differential expression is statistically significant are marked in red (FDR < 0.05 and |log2FC| > 1). (CDE) The expression of the HSP70 chaperone HSPA1A/HSPA1B was analyzed by qPCR in cells expressing the indicated ASPA variants. The HSP70 expression normalized to that in cells expressing WT ASPA is plotted (C) vs. the abundance score determined by VAMP-seq, (D) vs. the GFP/mCherry ratios determined in low throughput, and (E) vs. the toxicity score.
